# Diversity and community structure of anaerobic gut fungi in the rumen of wild and domesticated herbivores

**DOI:** 10.1101/2023.08.29.555426

**Authors:** Casey H. Meili, Moustafa A. TagElDein, Adrienne L. Jones, Christina D. Moon, Catherine Andrews, Michelle R. Kirk, Peter H. Janssen, Carl J. Yeoman, Savannah Grace, Joanna-Lynn C. Borgogna, Andrew P. Foote, Yosra I. Nagy, Mona T. Kashef, Aymen S. Yassin, Mostafa S. Elshahed, Noha H. Youssef

## Abstract

The rumen houses a diverse community that plays a major role in the digestion process in ruminants. Anaerobic gut fungi (AGF) are key contributors to plant digestion in the rumen. Here, we present a global amplicon-based survey of the rumen mycobiome by examining 206 samples from 15 animal species, 15 countries and six continents. The rumen mycobiome was highly diverse, with 81 out of 88 currently recognized AGF genera or candidate genera identified. However, only six genera (*Neocallimastix, Orpinomyces, Caecomyces, Cyllamyces,* NY9, and *Piromyces*) were present at > 4% relative abundance. AGF diversity was higher in members of the families *Antilocapridae* and *Cervidae* compared to *Bovidae*. Community structure analysis identified a pattern of phylosymbiosis, where host family (10% of total variance) and species (13.5%) partially explained the rumen mycobiome composition. Domestication (11.14%) and biogeography (14.1%) also partially explained AGF community structure, although sampling limitation, geographic range restrictions, and direct association between domestication status and host species hindered accurate elucidation of the relative contribution of each factor. Pairwise comparison of rumen versus fecal samples obtained from the same subject (n=13) demonstrated greater diversity and inter-sample variability in rumen over fecal samples. The genera *Neocallimastix* and *Orpinomyces* were present in higher abundance in rumen samples, while *Cyllamyces* and *Caecomyces* were enriched in fecal samples. Comparative analysis of global rumen and feces datasets revealed a similar pattern. Our results provide a global view of AGF community in the rumen and identify patterns of AGF variability between rumen and feces in herbivores tract.

**Importance:** Ruminants are highly successful and economically important mammalian suborder. Ruminants are herbivores that digest plant material with the aid of microorganisms residing in their GI tract. The rumen compartment represents the most important location where microbially-mediated plant digestion occurs in ruminants, and is known to house a bewildering array of microbial diversity. An important component of the rumen microbiome is the anaerobic gut fungi, members of the phylum Neocallimastigomycota. So far, studies examining AGF diversity have mostly employed fecal samples, and little is currently known regarding the identity of AGF residing in the rumen compartment, factors that impact the observed patterns of diversity and community structure of AGF in the rumen, and how AGF communities in the rumen compare to AGF communities in feces. Here, we examined the rumen AGF diversity using amplicon-based surveys targeting a wide range of wild and domesticated ruminants (n=206, 15 different animal species) obtained from 15 different countries. Our results demonstrate that while highly diverse, no new AGF genera were identified in the rumen mycobiome samples examined. Our analysis also indicate that animal host phylogeny plays a more important role in shaping AGF diversity in the rumen, compared to biogeography and domestication status. Finally, we demonstrate that a greater level of diversity and higher inter-sample variability was observed in rumen compared to fecal samples, with two genera (*Neocallimastix* and *Orpinomyces*) present in higher abundance in rumen samples, and two others (*Cyllamyces* and *Caecomyces*) enriched in fecal samples. Our results provide a global view of the identity, diversity, and community structure of AGF in ruminants, elucidate factors impacting diversity and community structure of the rumen mycobiome, and identify patterns of AGF community variability between the rumen and feces in the herbivorous GIT tract.

## Introduction

Ruminants (suborder *Ruminantia*) are one of the most diverse and prevalent groups of extant mammalian herbivores. The global population of domesticated ruminants is estimated at ∼3.75 billion, and that of wild ruminants is upwards of 75 million animals (1). Suborder *Ruminantia* includes six families: *Antilocapridae*, *Bovidae*, *Cervidae*, *Giraffidae*, *Moschidae*, and *Tragulidae* (2). The most species-rich family is *Bovidae* with >140 species (3), many of which are important livestock animals (e.g., cattle, goats, sheep) (1).

Ruminants are highly efficient in digesting high-fiber feeds and forages. This is primarily due to the ability to ferment feed in an anaerobic pregastric chamber (the rumen) which enables a specialized microbiome-mediated plant biomass degradation and fermentation to end products that are important energy sources for the hose (4). Rumination, the process by which animals regurgitate and masticate previously swallowed plant material, allows further physical breakdown of partially digested feed, enhancing rumen microbial activity.

The rumen microbial community encompasses bacteria, archaea, protozoa, and fungi (5). The fungal component, the anaerobic gut fungi (AGF), belongs to the phylum *Neocallimastigomycota*. They play a key role in the plant biomass degradation process. AGF hyphae efficiently penetrate plant biomass and mechanically disrupt plant cell walls (5), as well as by producing a wide array of carbohydrate-active enzymes (CAZymes) that are crucial for plant cell wall degradation (6–8). Indeed, several field studies have demonstrated the important contributions of AGF to biomass degradation in ruminants (7, 9, 10).

Surprisingly, in contrast to the wealth of information on the rumen bacterial and archaeal communities, little information is currently available on the resident rumen AGF community in the rumen of mammalian herbivores. The bulk of culture-independent and culture-based AGF characterization studies have been conducted on fecal, rather than rumen, samples due to the relative ease of sampling (11–14), and only a few studies to date have reported on AGF communities in rumen samples, all of which were limited in scope, examining few subjects and host species (15–19). AGF communities residing in the rumen could differ from those encountered in fecal samples due to possible selection and modification when passing through various partitions of the forestomach system (rumen, omasum, and abomasum) owing to the large difference in pH between these compartments (e.g., 6-6.4 in the rumen, 5.5-6.5 in the omasum, and 1.5-3 in the abomasum in cattle) before reaching the circumneutral small intestine. In addition, a fraction of fermentation occurs in the intestine (around 10%, (20)); and the potential AGF presence, origin, identity, load, and relative contribution to the AGF community encountered in fecal samples is currently unknown. Finally, distinct bacterial and archaeal communities colonize various locations in the GI tract of ruminants (21, 22), and these communities could differentially impact and modulate AGF diversity, load, and community composition through antagonistic, synergistic, or mutualistic relationships, as suggested in defined cocultures (23).

Because the rumen is the main site for feed digestion and absorption, studying the AGF community there provides insights into the taxa involved in active plant biomass degradation in ruminants. A detailed understanding of the AGF diversity and community structure in the rumen is key for devising strategies for community modulation, manipulation, and augmentation to improve the host’s overall health and feed efficiency. As well, the yet-unexamined rumen could represent a source for novel, hitherto uncharacterized AGF taxa that are selectively lost during feed passage from the rumen to the lower GI tract. To fill this knowledge gap, we characterized the AGF communities of a global collection of rumen samples (n=206) belonging to fifteen different species from three ruminant families using a culture-independent amplicon sequencing approach. In addition to the dataset’s broad host and geographic distribution, enabling biogeographic-based comparisons, it also allows comparison of domesticated (n=180) compared to wild (n=26) hosts, providing a unique opportunity to examine the effect of domestication on the rumen AGF community. Further, the rumen AGF community was compared to fecal AGF datasets obtained in a recent similar global survey. Finally, a direct pairwise comparison of AGF communities in the rumen and fecal samples simultaneously obtained from the same animal was conducted on a subset of animals. Our results provide a global view of the identity, diversity, and community structure patterns of AGF in the mammalian rumen and elucidate the role of phylogeny, biogeography, and domestication in structuring AGF communities. Further, rumen versus feces community comparisons suggest that while similar ecological and evolutionary factors impact the AGF community in both locations, distinct differences in the identity, diversity, and community structure patterns exist between both locations. We posit that such differences are driven by AGF acquisition routes, and the selection process associated with the transition of AGF from the pre-gastric rumen to the intestine.

## Materials and Methods

### Samples

A total of 206 rumen samples from 15 countries and six continents were obtained (Table S1, Figure 1a). Samples include representatives from three ruminant families (*Antilocapridae, Bovidae,* and *Cervidae*) and 15 different domesticated and wild animal species (Figure 1 b, Table S1). Many of these samples were obtained as part of a prior global rumen census (GRC) survey of the bacterial, archaeal, and protozoal communities in the rumen (24) (Table S1). Other samples were obtained from wild ruminants through collaboration with hunters at the state of Montana (n=27) (Table S1). A third fraction of samples (n=23) were collected from three slaughterhouses (Cairo, Giza and Menya) in Egypt with the help of trained technicians. The GITs of the slaughtered animals were separated on a clean bench and then sectioned with a knife. Rumen content solids were transferred in sterile labeled 50 ml Falcon tubes. The interval between the animal death and the sample collection did not exceed 30 min. The falcon tubes were stored at −20°C till DNA extraction (Table S1). Finally, samples were also obtained from an animal housed at the Oklahoma State University Department of Animal Sciences through gastric tube insertion (n=1, Table S1). All animal ethics approvals for rumen sampling from the GRC survey were obtained as outlined in (24). All hunters in Montana had the necessary hunting permits and harvested animals using legal methods. Sample collection and handling in Egyptian samples was approved by the Committee for Safe Handling and Disposal of Chemical and Biological Materials, Faculty of Pharmacy Cairo University # MI3011, June 2021. The sampling procedure at Oklahoma State University was reviewed and approved by the Oklahoma State University Institutional Animal Care and Use Committee (Protocol #21-03).

**Figure 1.**
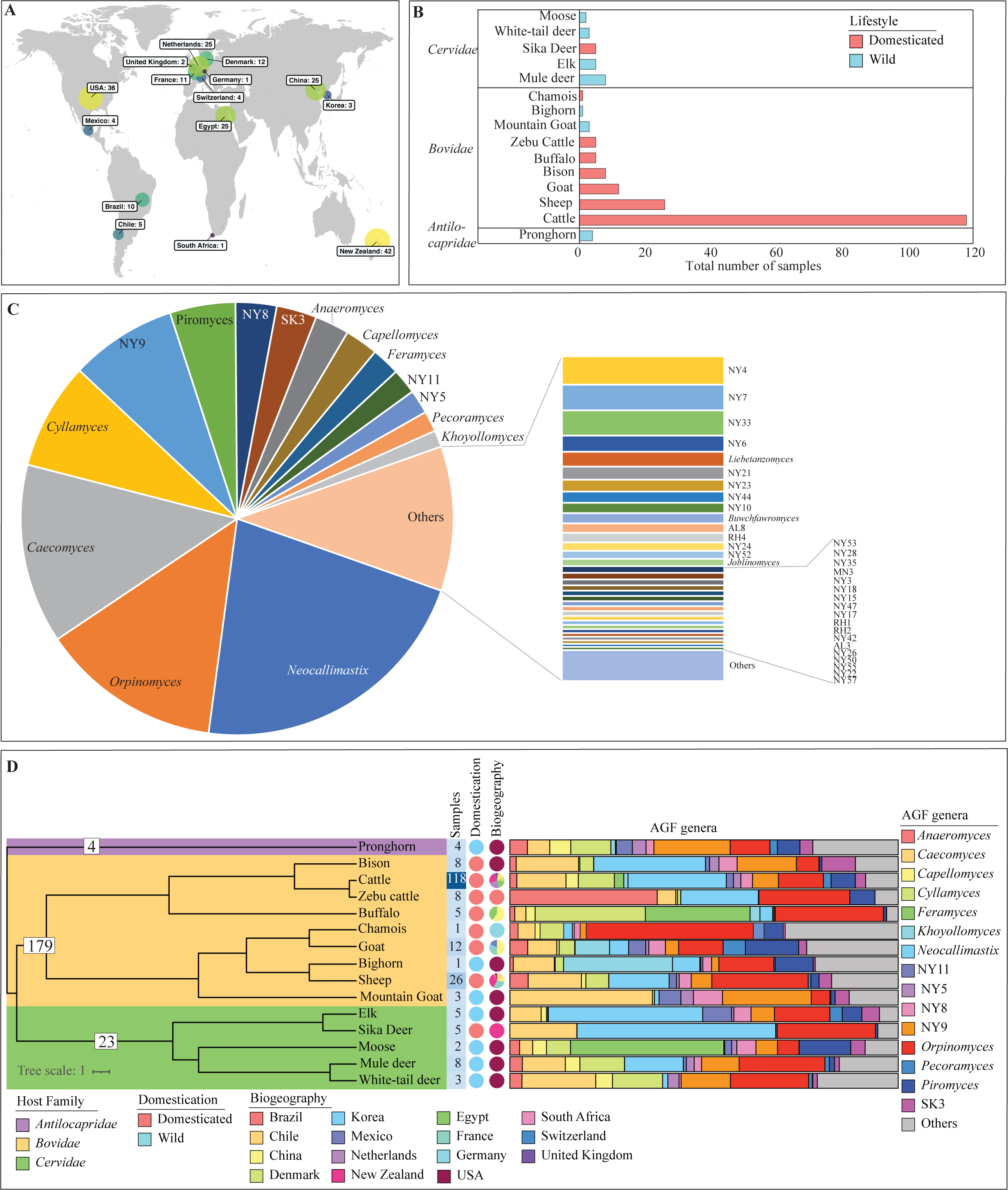
AGF diversity patterns in the rumen mycobiome. (A) A map showing the biogeographical origin of all samples (n=206) analyzed in this study. (B) Bar plots showing the number of samples belonging to each animal species. Animals are ordered by family. Domestication status is color-coded (domesticated in peach and wild in cyan). (C) Pie chart showing the overall composition of AGF genera encountered in the entire (1.86 million sequence) dataset. Genera present in >1% relative abundance are named on the pie chart, genera with relative abundances 0.5-1% are named in the bar chart to the right, and genera present in <0.5% relative abundance are collectively referred to as “Others”. (D) AGF community composition in the animal species studied. The phylogenetic tree downloaded from timetree.org shows the relationship between the 15 species sampled. Species are color-coded by their family and the total number of samples belonging to each of the three families is shown at each corresponding node. “Samples” refer to the number of samples belonging to each animal species and is shown to the right of the tree as a heatmap with the exact numbers displayed. “Domestication” refers to the domestication status (domesticated in peach and wild in cyan) and is shown as a pie chart to the right of the heatmap. “Biogeography” shows the distribution of the number of samples from different geographical regions and is shown as a pie chart to the right of “Domestication”. Countries are color-coded as shown in the figure key. The AGF community composition for each animal species is shown to the right as colored columns corresponding to the legend key. Genera with a total abundance of >1% are shown, while all other genera are grouped as “Others”.

### DNA extraction, amplification, and sequencing

DNA from the GRC rumen samples was extracted as described previously (24) and stored at −80^0^C, then stabilized using GenTegra-DNA protectant (GenTegra, Pleasanton CA, USA) to minimize DNA degradation during shipping from New Zealand to Oklahoma State University. For all other samples, whole (i.e., solid and liquid) rumen samples were collected, frozen, and transferred to the laboratory where they were promptly stored at −80°C. DNA was extracted from all samples using the DNeasy Plant Pro Kit (Qiagen, Germantown, MD, USA) according to the manufacturer’s instructions. PCR amplification reactions, amplicon clean-up, quantification, index and adaptor ligation, and multiplexing were conducted in a single laboratory (Oklahoma State University, Stillwater, OK, USA) to eliminate inter-laboratory variability. The procedure, previously outlined in detail in reference (13), involved amplification of a ∼370 bp of the second variable region of the large ribosomal subunit (D2-LSU) using primers AGF-LSU-EnvS primer pair (AGF-LSU-EnvS For: 5’-GCGTTTRRCACCASTGTTGTT-3’, AGF-LSU-EnvS Rev: 5’-GTCAACATCCTAAGYGTAGGTA-3’). Pooled libraries were sequenced at the University of Oklahoma Clinical Genomics Facility (Oklahoma City, OK, USA) using an Illumina MiSeq platform as previously described (13).

### Sequence processing and phylogenetic placement

Protocols for read assembly and sequence quality trimming, as well as procedures for calculating thresholds for species and genus delineation and genus-level assignments were conducted as described previously (13). Assignment of sequences to AGF genera was conducted using a two-tier approach for genus-level phylogenetic placement and thresholds as described previously (11, 13, 25).

### Diversity and community structure assessment

The relationship between host identity and AGF diversity and community structure was examined across animal host families and species for animals with at least four samples at each of these levels. This included the three animal host families, and the animal species pronghorn (n=4), cattle (n=116), sheep (n=26), goat (n=14), American bison (n=8), water buffalo (n=5), zebu cattle (n=5), mule deer (n=8), sika deer (n=5), and elk (n=5). The effect of biogeography was examined by clustering samples by country and only including countries with at least 4 samples. This included samples from New Zealand (n=42), USA (n=36), China (n=25), Egypt (n=25), Netherlands (n=25), Denmark (n=12), France (n=11), Brazil (n=10), Chile (n=5), Mexico (n=4), and Switzerland (n=4). However, due to the uneven distribution of animals across locations, we also re-analyzed the effects of biogeography across the same animal species. Only cattle and sheep were considered for this comparison as they had enough representation (at least 4 samples) across different countries (Brazil, China, Denmark, Egypt, Netherlands, New Zealand, and Switzerland for cattle, and Chile, France, and New Zealand for sheep). The effect of domestication status was also examined by clustering animals into wild or domesticated categories. However, domestication status could also be conflated with host phylogeny due to unequal representation of wild animals from the families *Cervidae*, and *Antilocapridae*, and domesticated animals in family *Bovidae*. Domestication status could also be conflated with biogeography, with all wild animals originating from samples obtained in the USA. To partially alleviate this issue, we also examined the effect of domestication status within members of the same family, comparing wild bighorn and mountain goat (n=4) versus all other domesticated *Bovidae* species, and domesticated sika deer (n=5) versus all other wild *Cervidae* species. Finally, domesticated versus wild comparison of members of the same animal genus was also conducted was possible (for domesticated members of the genus *Cervus* (sika deer, *C. nippon*, n=5) versus wild (elk, *C. canadensis*, n=5)).

Diversity indices (Shannon, Simpson, and Inverse Simpson) were calculated using the estimate_richness command in the phyloseq R package, and the effect of factors (host family, species, domestication status, and biogeography) on alpha diversity was calculated using the aov command in R. The TukeyHSD command in R was used for multiple comparisons of means on the ANOVA results for all pairwise comparisons.

For community structure analysis, weighted Unifrac was calculated using the distance command in the phyloseq R package. Pairwise values were used to construct PCoA ordination plots using the commands ordinate and plot_ordination in phyloseq R package. To elucidate factors significantly impacting community structure, PERMANOVA tests were run using the command adonis in vegan R package. Percentage variance explained by each factor was calculated as the percentage of the sum of squares of each factor to the total sum of squares, and F-statistics p-values were used to examine the significance of the effect.

Multiple regression of matrices (MRM), Mantel tests for matrices correlations, and Procrustes rotation were also utilized to further quantify factors that could explain the divergence in AGF communities. MRM and Mantel tests were conducted by comparing a Gower-transformed matrix of each host factor (host family, species, domestication status, and biogeography) to the weighted Unifrac beta diversity dissimilarity matrix (calculated as detailed above) using the MRM and Mantel commands in the ecodist R package. Gower transformation of host factor matrices was conducted using the daisy command in the cluster R package. The protest command in the vegan R package was utilized for Procrustes rotation calculations. P-values, and coefficients (R^2^ regression coefficients from MRM analysis, Spearman correlation coefficients from Mantel tests, and symmetric orthogonal Procrustes statistic from Procrustes analysis) were examined to determine the significance, and importance of factors, respectively, in shaping the AGF community.

To identify specific animal host-fungal associations, LIPA (Local Indicator of Phylogenetic Association) was employed using the lipaMoran command in the phylosignal R package. For genera with significant associations (p-value <0.05), we calculated the average LIPA value for each animal species. Only genera with >1% relative abundance in the entire dataset (n=15) were examined. We considered average LIPA values in the range of 0.2-0.4 to represent weak associations, in the range of 0.4-1 to represent moderate associations, and above 1 to represent strong associations.

### AGF diversity in rumen versus fecal samples

For a subset of animals, (12 cattle and one water buffalo, Table S1) fecal samples were obtained as the same time as rumen samples. One cattle subject was housed at the Oklahoma State Animal Sciences Department. Rumen samples were obtained via gastric tubing, as described above, and the first fecal sample deposited after rumen collection was obtained. For the remaining animals (11 cows and 1 buffalo), subjects were slaughtered as part of the slaughterhouse operations, and rumen and fecal samples were directly obtained post-slaughter. DNA extraction, amplification, and sequencing of fecal samples were conducted following the same procedures for rumen samples outlined above.

As well, we sought to evaluate whether the observed patterns from pairwise comparison of samples obtained from the same animal could be extrapolated to larger datasets where fecal and rumen samples were obtained from different animals. To this end, we compared the community structure of cattle rumen (n=116) and fecal (n=178) AGF communities using all cattle rumen samples obtained in this study and cattle fecal samples obtained in a recent global survey of the AGF mycobiome (13).

For both rumen versus feces datasets obtained from the same animal (n=26), and global cattle rumen versus feces (n=294), DPCoA plots were calculated using the plot_ordination command in the phyloseq R package. In addition, metastats (26) in mothur was used to identify genera differentially abundant in rumen versus feces samples.

### Sequence and data deposition

Illumina amplicon reads generated in this study have been deposited in GenBank SRA under BioProject accession number PRJNA1008183 and Biosample accessions numbers SAMN37111842-SAMN37112060.

## Results

### Rumen AGF community overview

Illumina sequencing of 206 different rumen samples generated 1.86 million (average=9029) high-quality AGF-affiliated D2-LSU sequences (Table S2). High coverage values (average 0.996, minimum 0.92, coverage higher than 0.98 in 197/206 samples) indicated that the majority of genus-level diversity was captured in all samples (Table S2). Phylogenetic analysis identified 81 of the 88 AGF genera currently described (22/22 cultured; 59/66 uncultured) (Table S2, Figure 1c, d), and no new AGF genera were identified in the dataset. While the majority of currently recognized AGF genera and candidate genera were encountered, only 15 genera were present at >1% abundance in the entire dataset (Figure 1c), and only seven genera (*Neocallimastix, Caecomyces, Orpinomyces, Caecomyces, Cyllamyces,* NY9, and *Piromyces*) were present at >4% abundance (Figure 1c). Relative abundance and occurrence for AGF genera were highly correlated (R^2^ = 0.571, Figure S1).

### Patterns of alpha diversity in the rumen mycobiome

Alpha diversity patterns were assessed using three different indices: Shannon, Simpson, and Inverse Simpson (Figure 2, Figure S2). Collectively, samples belonging to the family *Bovidae* are less diverse when compared to members of the families *Cervidae* and *Antilocapridae* (Figure 2a, Figure S2a-b). Within specific animal species, the AGF community in the rumen of pronghorn (family *Antilocapridae*), mule deer and elk (family *Cervidae*) were the most diverse, while goat (family *Bovidae)* harbored the least diverse community. Pairwise differences in estimates of AGF community alpha diversity were significant between host species belonging to different families, e.g. pronghorn (*Antilocapridae*) versus goat (*Bovidae*), mule deer (*Cervidae*) versus goat and cattle (*Bovidae*), and sika deer (*Cervidae*) versus goat (*Bovidae*), as well as within few pairs of species in the family Bovidae, e.g. American Bison versus goat, sheep versus goat, and sheep versus cattle (Figure 2a, Figure S2a-b).

**Figure 2.**
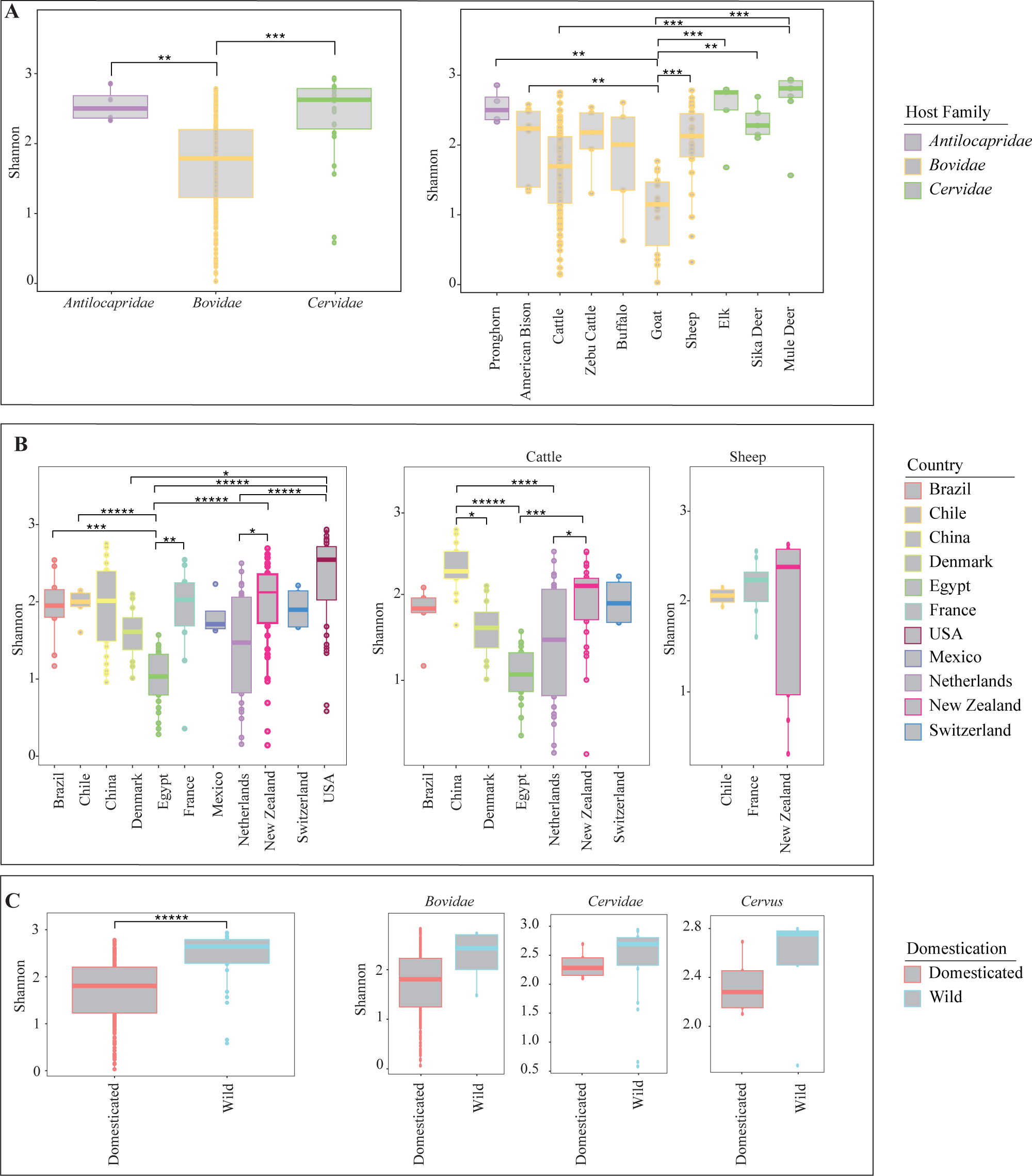
AGF alpha diversity in the rumen mycobiome. (A) Box and whisker plots showing the distribution of Shannon diversity measure for different animal families (left), and animal species (right) with four or more samples. (B) Box and whisker plots showing the distribution of Shannon diversity measure for animals from different biogeographical locations (only countries with at least 4 samples are shown). Results are shown for the total dataset (left), and for only cattle (middle), or only sheep (right). (C) Box and whisker plots showing the distribution of Shannon diversity measure for domesticated versus wild animals. Results are shown for the total dataset (left), and for animals belonging to the families *Bovidae*, *Cervidae*, or the genus *Cervus* as depicted on top of each figure. Results for two-tailed ANOVA followed by Tukey for pairwise animal family and animal species comparisons are shown on top of the box plots only for significant comparisons. *, 0.01 < p < 0.05; **, p < 0.01; ***, p < 0.001; ****, p < 0.0001; *****, p < 0.00001.

In addition to host identity, multiple pairwise significant differences were observed between alpha diversity patterns of samples when grouped by the country of origin (Figure 2b). However, since biogeographic patterns could be a reflection of unequal distribution of hosts species across locations as described above, we also examined differences in alpha diversity between the same animal species from different countries. Only cattle and sheep had adequate samples (at least 4) across different countries (Brazil, China, Denmark, Egypt, Netherlands, New Zealand, and Switzerland for cattle, and Chile, France, and New Zealand for sheep) to enable such analysis. We identified significant differences in alpha diversity based on country of origin in cattle, but not sheep (Figure 2b, S2c-d). Finally, a comparison of alpha diversity estimates between samples from domesticated versus wild animals revealed higher levels of diversity in wild compared to domesticated hosts (Figure 2c), although such differences were not significant when restricting the analysis to animal hosts within the same host family or genus (Figure 2c).

### Rumen mycobiome community structure

AGF community structure analysis indicated a significant role for host phylogeny in shaping AGF rumen mycobiome at the family (p=0.001) and the species (p=0.001) levels; although such factors explained only 10.0% and 13.5% of variance, respectively (Figure 3a). Similarly, domestication status was a significant factor (p=0.001) in explaining 11.1% of the AGF rumen mycobiome variance when using the entire dataset (Figure 3b), as well as when subsetting samples belonging to the genus *Cervus* (p=0.01), but not when subsetting samples belonging to the families *Bovidae* and *Cervidae* (Figure 3b). Finally, biogeography was also significantly associated with the AGF rumen mycobiome community, explaining 14.1% of the community variance using the entire dataset level (Figure 3c). A significant effect of biogeography on AGF community structure was also observed when restricting the analysis to a single host species (cattle and sheep, Figure 3c).

**Figure 3.**
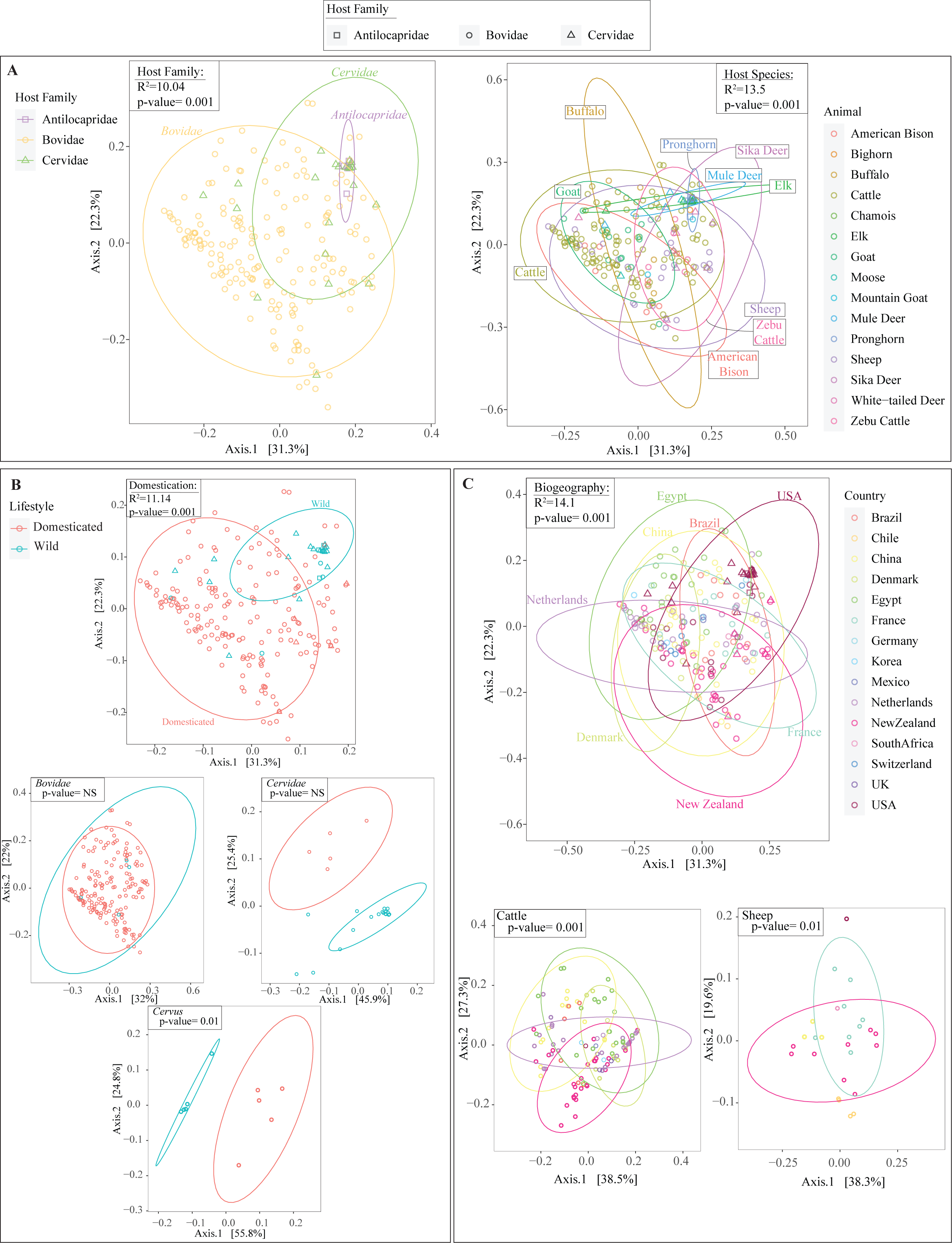
Patterns of AGF beta diversity in the rumen mycobiome. Principal coordinate analysis (PCoA) ordination plots based on AGF community structure in the 206 samples studied here constructed using the phylogenetic similarity based Unifrac weighted. The shape represents the animal family as shown on top. The % variance explained by the first two axes are displayed on the axes, and ellipses encompassing 95% of variance are displayed. In (A), samples and ellipses are color coded by animal family (left), or animal species (right). In (B), samples and ellipses are color coded by animal domestication status when using the total dataset (top), or only for animals belonging to the families *Bovidae* (middle left), *Cervidae* (middle right), or the genus *Cervus* (bottom). In (C), samples and ellipses are color coded by animal biogeography when using the total dataset (top), or only for cattle (bottom left), or sheep (bottom right). PERMANOVA results for partitioning the dissimilarity by variation sources (animal family, animal species, domestication status, and country) is shown for each plot. R^2^ refers to percentage variance explained by each factor (calculated as the percentage of the sum of squares of each factor to the total sum of squares), while p-value refers to the F-statistics p-value.

In addition to PERMANOVA, multiple regression of matrices (MRM), Mantel tests for matrices correlations, and Procrustes rotation were utilized to quantify factors that could explain the divergence in AGF communities. Results of matrices correlation using each of the three methods confirmed the importance of animal host species, family, biogeography, and domestication status in explaining the AGF community structure (Figure S3). Finally, to identify whether specific AGF genera are associated with specific animal hosts and to quantify the strength of such associations, we employed LIPA analysis. For the 15 genera encountered at >1% abundance, LIPA analysis identified 17 (5 weak, 4 moderate, and 8 strong) AGF genus-animal host associations (Figure S4).

### Rumen-feces mycobiome comparison

Direct comparisons were made of rumen and fecal samples obtained simultaneously from 12 cows and one water buffalo sample. Within this relatively limited dataset, clear differences were observed in the AGF community composition (Figure 4a), alpha diversity (Figure 4b), and community structure (Figure 4c). Fecal samples AGF communities were significantly less diverse than those from rumen samples (Figure 4b). DPCoA ordination plots using weighted Unifrac demonstrated clear clustering of rumen and fecal communities, with sampling location (rumen versus feces) explaining 51.66% of community variance. The level of variability within each sampling location was quantified by measuring the variability in Euclidean distances of samples from each sampling location to their corresponding group centroid (the centroid of the 95% ellipses shown in Figure 4c). A significantly greater level of variability was observed in rumen versus feces samples. Further, DPCoA showed selective enrichment of specific AGF taxa for each sampling location with the genera *Neocallimastix*, and *Orpinomyces* selectively enriched in the rumen, and the genera *Caecomyces* and *Cyllamyces* selectively enriched in fecal samples. Metastats confirmed significance of these specific genera selective enrichment.

**Figure 4.**
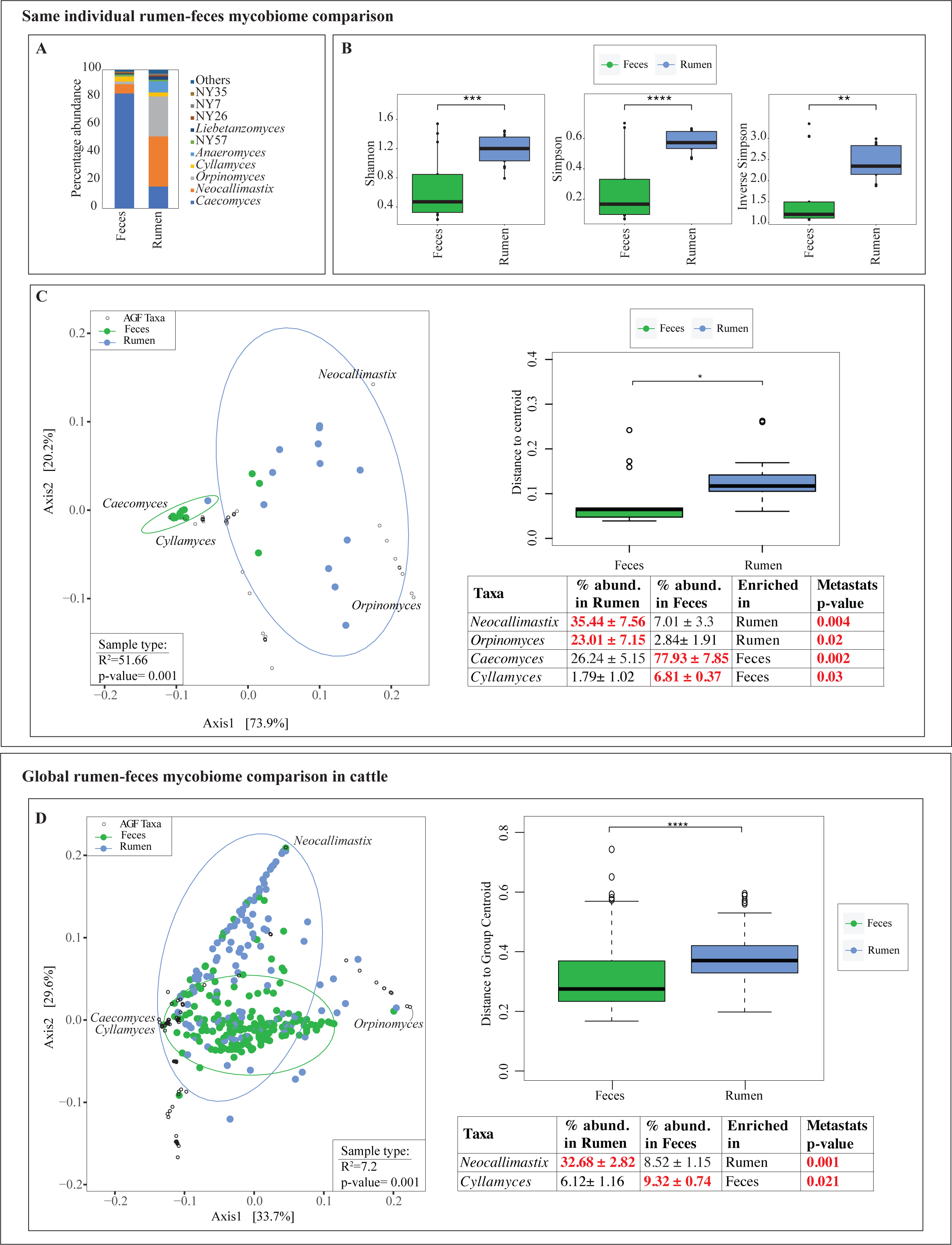
Rumen-feces mycobiome comparison. (A-C) Same individual rumen-feces mycobiome comparison conducted on 13 animal subjects (12 cattle, and 1 buffalo). (A) Collective AGF community composition for each sampling location. Genera with >1% total abundance are color coded as shown to the right. All other genera are grouped as “others”. (B) AGF alpha diversity patterns in the 13 rumen-versus-feces samples. Box and whisker plots show the distribution of Shannon (left), Simpson (middle), and Inverse Simpson (right) diversity indices in the two sampling locations. (C) Double principal coordinate analysis (DPCoA) biplot based on the phylogenetic similarity-based index weighted Unifrac showing the community structure in the 13 rumen and 13 feces samples. The % variance explained by the first two axes is displayed on the axes, and ellipses encompassing 95% of variance are displayed. The samples and ellipses are color-coded by sampling location (rumen, blue; feces, green). AGF genera are shown as smaller black empty circles and the four AGF genera with selective enrichment in either sampling locations are labeled. PERMANOVA results are shown in the bottom left corner of the plot, where R^2^ refers to percentage variance explained by the sampling location (calculated as the percentage of the sum of squares of each factor to the total sum of squares), while p-value refers to the F-statistics p-value. To the right of the DPCoA plot, the level of variability between samples from the same sampling location is shown as box and whisker plots for the distribution of DPCoA ordination distance of each sample to its group centroid, and results for two-tailed ANOVA is shown on top of the box plots: *, 0.01 < p < 0.05. Results of metastats for these four genera are shown in the table. For each taxon, the average and standard deviations of abundance is shown for the rumen versus feces, followed by the sampling location where the taxon was identified as significantly differentially abundant, and the metastats p-value. (D) Global cattle rumen-feces mycobiome comparison conducted on the cattle rumen samples analyzed in this study (n=116) and 178 cattle fecal samples obtained in a recent global survey of the AGF mycobiome (13). Double principal coordinate analysis (DPCoA) biplot based on the phylogenetic similarity-based index weighted Unifrac showing the community structure in the 116 rumen and 178 feces samples. The % variance explained by the first two axes is displayed on the axes, and ellipses encompassing 95% of variance are displayed. The samples and ellipses are color-coded by sampling location (rumen, blue; feces, green). The same four AGF genera identified as selectively enriched in either sampling locations in (C) with are labeled. PERMANOVA results are shown in the bottom left corner of the plot, where R^2^ refers to the percent variance explained by the sampling location, while p-value refers to the F-statistics p-value. The level of variability between samples from the same sampling location is shown as box and whisker plots for the distribution of DPCoA ordination distance of each sample to its group centroid, and results for two-tailed ANOVA is shown on top of the box plots: ****, p < 0.0001. Results of metastats for the two genera with significant differential abundance are shown.

We sought to evaluate whether the observed patterns of AGF genera selective enrichment in rumen versus feces could be extrapolated to a more global level on datasets where fecal and rumen samples were obtained from different animals. We, thus, compared the community structure of cattle rumen and fecal AGF communities using the cattle rumen samples analyzed in this study (n=116) and 178 cattle fecal samples obtained in a recent global survey of the AGF mycobiome (13). For this larger dataset (Figure 4d), DPCoA ordination plots using weighted Unifrac showed significant clustering with sampling location (rumen versus feces), albeit explaining only a minor fraction of the community variance (7.2%). Similar to the smaller dataset, a significantly greater degree of variability was observed in rumen versus feces samples (Figure 4d). While DPCoA showed a very similar pattern for the four AGF taxa identified above (*Neocallimastix*, and *Orpinomyces* clustering close to rumen samples, and the genera *Caecomyces* and *Cyllamyces* clustering close to fecal samples) (Figure 4d), Metastats analysis only confirmed significance of the selective enrichment of *Neocallimastix* in cattle rumen and *Cyllamyces* in cattle feces (Figure 4d).

## Discussion

We present a detailed assessment of the rumen compartment mycobiome in 206 mammalian herbivores using a culture-independent amplicon-based survey. Our results highlight the high level of overall gamma diversity within the global rumen mycobiome, with 81 out of the 88 currently reported AGF genera identified (Figure 1c). The AGF rumen mycobiome community composition displayed a pattern where a relatively limited number of genera were ubiquitous (occurring in >50% of the samples) and abundant (representing a large fraction of the AGF community when encountered) (Figures 1d, S1). The remaining AGF genera displayed lower levels of occurrence and relative abundance (Figure S1). Such a pattern of high diversity and predominance of few genera is consistent with prior surveys of AGF in fecal samples of herbivores (13, 27). The rationale behind the existence and maintenance of perpetually rare genera within the herbivorous gut has previously been debated, and is potentially attributed to their superior survival capabilities or probable role played under specific conditions not adequately captured in the current sampling schema (e.g., younger age, stress, specific types of feed) (13). Significantly, our analysis failed to identify novel AGF genera beyond those previously observed in prior feces-based surveys (Figure 1c-d, (11-14, 18)), hence refuting the preposition that rumen samples could represent a significant reservoir of novel, hitherto undescribed AGF diversity. This does not preclude novel AGF diversity in wild ruminants with specialized diets or feeding behaviors, like reindeer feeding on lichen or browsing ruminants in tropical forests, but widespread novelty seems unlikely.

Beyond documenting the occurrence and relative abundance (Figure 1) of AGF taxa, we examined patterns of their diversity and community structure in the rumen mycobiome and attempted to elucidate the role of and interplay between various factors in shaping the observed patterns. Our results document statistically significant differences in levels of diversity (Figures 2, S2) and community structure (Figure 3) patterns between various families and host species, suggesting a pattern of phylosymbiosis, where host phylogenetic affiliation plays a role in shaping the AGF community. As well, LIPA analysis (Figure S4) has shown few specific pairwise AGF genus-animal species associations (e.g., goat with *Caecomyces*, *Orpinomyces*, *Cyllamyces*, and *Neocallimastix*, buffalo with *Orpinomyces*, American bison with NY9, pronghorn, elk, and mule deer with *Khoyollomyces*). Interestingly, recent work on the fecal AGF mycobiome has also identified patterns of phylosymbiosis, with specific LIPA preferences largely concordant in most animals shared between the two datasets (e.g. buffalo with *Orpinomyces*, elk and mule deer with *Khoyollomyces*).

It is important to note that quantitative assessment of the role of host identity in explaining rumen AGF community structure (PERMANOVA, and multivariate matrices comparisons using MRM, Mantel, and Procrustes) indicates that, host species/family could explain only a relatively small fraction of the observed variance (Figures 3, S3). This indicates that additional factors such as domestication status, and biogeography could possibly play an additional role in shaping the rumen AGF community. Domesticated animals typically receive a less diverse, more frequent and homogenous dietary regiment that is often grain-rich. This is in stark contrast to wild herbivores that browse or graze on a more heterogenous diet with a feeding regiment controlled by resource availability and predation risk. Our results indicate a higher level of AGF alpha diversity in wild animals (Figure 2c, S2e-f), and a significant role (F-statistic R^2^=11.14%, p-value=0.001) for domestication status in shaping AGF community (Figure 3b, S3). As such, we posit that more variable feed types and non-monotonous feeding patterns in wild herbivores could lead to enrichment and co-existence of a more diverse AGF community suited to a more stochastic feeding regiment. However, it is important to note that all animal species examined were either exclusively wild or domesticated, leading to a potential conflation of both factors (animal species and domestication status) as drivers of AGF diversity and community structure. We attempted to partially control for the conflation of both factors by re-analyzing the impact of domestication on AGF diversity and community structure on subsets of the datasets comprised of animals from the same family (families *Bovidae* and *Cervidae*, Figures 2c, S2e-f, 3b) or species (genus *Cervus*, Figures 2c, S2e-f, 3b). The results hint at a potential (albeit not significant) role for domestication towards lower diversity and selection of taxa, as evidenced by differences between closely related wild and domesticated animals. Nevertheless, only a highly controlled experiment, where wild and domesticated subjects belonging to the same animal species from the same location are compared could conclusively disentangle both factors, e.g., capturing, rearing, and sampling white-tailed deer species in a domesticated setting and comparing their AGF community to wild deer from the same region.

Biogeography could be an additional factor impacting AGF diversity and community structure, as previously postulated for rumen bacterial and archaeal communities (24). Our results show that biogeography could play a role in shaping AGF diversity (Figures 2b, S2c-d) and community structure (Figure 3c). However, similar to domestication status, the result of biogeographic-based assessments could be skewed by the over-representation of specific animal species in certain locations. We attempted to partly disentangle host and biogeography by reanalyzing subsets constituting the same animal species from different locations. Our results suggest a role for biogeography in shaping AGF diversity in cattle. It is interesting to note that a similar observation was also discerned in a recent global dataset of fecal samples (13). The role of biogeography in shaping the AGF community could be driven by variability in cattle breed anatomic characteristics, feeding regiment, and rearing conditions between two locations.

Prior studies on AGF diversity in ruminants have largely been conducted on fecal, rather than rumen samples (11-14, 18). The lack of studies on AGF communities in the rumen was largely hampered by methodological limitations. Collection of rumen samples requires surgical fistulation or gastric tubing, processes that could be conducted in research settings, but are largely unfeasible for a broad sampling of herds in farming and ranching settings (28, 29). In wild ruminants, such an approach is not feasible, except in extremely rare conditions, where domestication of a naturally wild host was achieved (30). Theoretically, differences in AGF community between rumen and feces could be driven by selection for or against specific AGF taxa when passing through various regions within the animal’s alimentary tract, enrichment of specific AGF genera involved in intestinal fermentation (20)), or interaction between AGF and the distinct bacterial and archaeal communities colonizing various location in the animal’s alimentary tract. Using a pairwise sampling scheme in 13 animal subjects, we sought to assess differences between AGF communities in rumen versus feces samples. We acknowledge the relatively limited number of replicates and restriction to mostly one species (*Bos taurus*) and hence the patterns obtained should be regarded as preliminary. Our analysis clearly demonstrated that the AGF community in rumen samples is significantly more diverse than feces (Figure 4b). As well, distinct differences in community structure were observed between rumen and feces samples, with a selective enrichment of the genera *Neocallimastix* and *Orpinomyces* in rumen sample and *Caecomyces* and *Cyllamyces* in fecal samples. The underlying reasons for the observed inhibition and enrichment trends are presently unclear, given our current rudimentary knowledge regarding fine differences in metabolic and physiological preferences between various AGF genera. Nevertheless, it is notable that the genera *Caecomyces* and *Cyllamyces* are the only known AGF exhibiting a bulbous rhizoidal growth pattern and appear to have a unique attachment/pressing on plants compared to filamentous rhizoids. This growth pattern, with a higher proportion of the fungal thallus protected within the plant biomass, compared to the more superficial external hyphal attachment pattern in filamentous genera could offer a better protection during rumen contents passage through the highly acidic abomasum to the intestine. As well, while all AGF appear to grow readily and specialize in attacking intact plant biomass, a differential preference or efficiency of some genera in attacking, penetrating, and colonizing intact plant biomass would confer a competitive advantage in the rumen, where intact plants are first acted upon by the animal’s microbiome. On the other hand, a greater affinity for oligomers, dimers, and monomers uptake could enrich specific genera in the colon, where available substrates are mostly soluble sugars rather than intact plant material. It is interesting to note that distinct differences in rumen versus fecal communities have also been observed in bacteria and archaea, where a similar pattern of lower diversity in feces was observed, as well as a distinct preference for fiber-degrading taxa (e.g. *Fibrobacter*) in rumen as opposed to sugar-degrading taxa (e.g. *Tenericutes*) in feces (31, 32).

## Acknowledgments

We thank the following members of the GRC project for contributing samples used in this study:

**Olubukola Ajike Isah**: Department of Animal Nutrition, Federal University of Agriculture, Abeokuta (FUNAAB), Nigeria.

**Jorge Avila-Stagno**: Facultad de Ciencias Veterinarias, Universidad de Concepción, Chillan, Chile.

**Kasper Dieho, Jan Dijkstra, and Andre Bannink**: Animal Nutrition Group, Wageningen University, 6700 AH Wageningen, The Netherlands.

**Fabian N. Fon**: Department of Agriculture, University of Zululand, KwaDlangezwa, Empangeni, 3886, South Africa.

**Hilario Mantovani**: Departamento de Microbiologia, Universidade Federal de Viçosa, Campus UFV, 36570-000 Viçosa, Minas Gerais, Brazil.

**Martin Fraga**: Departamento de Microbiología, Instituto de Investigaciones Biológicas Clemente Estable, Av. Italia 3318, CP 11600, Montevideo, Uruguay.

**Francisco E. Franco:** VITA Marangani, Universidad Nacional Mayor de San Marcos, Lima, Perú.

**Chris Friedman**: Ministry for Primary Industries Verification Services Hawkes Bay, Silver Fern Farms—Pacific, Whakatu, Hastings, New Zealand.

**Arjan Jonker and Cesar S. Pinares-Patino**: AgResearch Limited, Grasslands Research Centre, Palmerston North 4442, New Zealand.

**Sophie Krizsan**: Department of Agricultural Research for Northern Sweden, Swedish University of Agricultural Sciences, SE-901 83 Umea, Sweden.

**Jan Lassen**: Department of Molecular Biology and Genetics, Aarhus University, DK-8830 Tjele, Denmark.

**Satoshi Koike and Yasuo Kobayashi**: Laboratory of Animal Nutrition, Research Faculty of Agriculture, Hokkaido University, N9W9, Kita-ku, Sapporo 060–8589, Japan.

**Sang Suk Lee and Lovelia L. Mamuad**: Department of Animal Science & Technology, Sunchon National University, Suncheon, Jeonnam 540–742, Korea.

**Cécile Martin, Diego Morgavi, and Milka Popova**: Institut National de la Recherche Agronomique, UMR1213 Herbivores, Saint-Genes-Champanelle, F-63122, France.

**Andreas Muenger**: Agroscope, Institute for Livestock Sciences ILS, CH-1725 Posieux, Switzerland.

**Camila Muñoz, Rodrigo de la Barra, María Eugenia Martínez, and Andrés M. Carvajal**: Instituto de Investigaciones Agropecuarias, INIA Remehue, Osorno, Región de Los Lagos, Chile.

**Mario A. Cobos-Peralta:** Colegio de Postgraduados, Institución de Ensenanza e Investigación en Ciensias Agrícolas, CP 56230, Montecillo, Mexico.

**Tasia Taxis**: Animal Science Genetics, University of Missouri-Columbia, Columbia, Missouri 65211, USA.

**Emilio M. Ungerfeld**: Instituto de Investigaciones Agropecuarias, INIA Carillanca, Temuco, Chile.

**Min Wang and Zhi Liang Tan**: Institute of Subtropical Agriculture, The Chinese Academy of Sciences, Changsha, China.

**Tianhai Yan**: Agri-Food and Biosciences Institute, Hillsborough, County Down BT26 6DR, Northern Ireland.

## References

1. Hackmann TJ, Spain JN. 2010. Invited review: ruminant ecology and evolution: perspectives useful to ruminant livestock research and production. J Dairy Sci 93:1320–34.

2. Decker JE, Pires JC, Conant GC, McKay SD, Heaton MP, Chen K, Cooper A, Vilkki J, Seabury CM, Caetano AR, Johnson GS, Brenneman RA, Hanotte O, Eggert LS, Wiener P, Kim JJ, Kim KS, Sonstegard TS, Van Tassell CP, Neibergs HL, McEwan JC, Brauning R, Coutinho LL, Babar ME, Wilson GA, McClure MC, Rolf MM, Kim J, Schnabel RD, Taylor JF. 2009. Resolving the evolution of extant and extinct ruminants with high-throughput phylogenomics. Proc Natl Acad Sci USA 106:18644–9.

3. Heller R, Frandsen P, Lorenzen ED, Siegismund HR. 2013. Are there really twice as many bovid species as we thought? Syst Biol 62:490–3.

4. Moraïs S, Mizrahi I. 2019. Islands in the stream: from individual to communal fiber degradation in the rumen ecosystem. FEMS Microbiol Rev 43:362–379.

5. Gruninger RJ, Puniya AK, Callaghan TM, Edwards JE, Youssef N, Dagar SS, Fliegerova K, Griffith GW, Forster R, Tsang A, McAllister T, Elshahed MS. 2014. Anaerobic fungi (phylum *Neocallimastigomycota*): Advances in understanding their taxonomy, life cycle, ecology, role and biotechnological potential. FEMS Microbiol Ecol 90:1–17.

6. Couger MB, Youssef NH, Struchtemeyer CG, Liggenstoffer AS, Elshahed MS. 2015. Transcriptomic analysis of lignocellulosic biomass degradation by the anaerobic fungal isolate *Orpinomyces* sp. strain C1A. Biotechnol Biofuels 8:208.

7. Hagen LH, Brooke CG, Shaw CA, Norbeck AD, Piao H, Arntzen MØ, Olson HM, Copeland A, Isern N, Shukla A, Roux S, Lombard V, Henrissat B, O’Malley MA, Grigoriev IV, Tringe SG, Mackie RI, Pasa-Tolic L, Pope PB, Hess M. 2021. Proteome specialization of anaerobic fungi during ruminal degradation of recalcitrant plant fiber. ISME J 15:421–434.

8. Youssef NH, Couger MB, Struchtemeyer CG, Liggenstoffer AS, Prade RA, Najar FZ, Atiyeh HK, Wilkins MR, Elshahed MS. 2013. The genome of the anaerobic fungus *Orpinomyces* sp. strain C1A reveals the unique evolutionary history of a remarkable plant biomass degrader. Appl Environ Microbiol 79:4620–34.

9. Elliott R, Ash AJ, Calderon-Cortes F, Norton BW, Bauchop T. 1987. The influence of anaerobic fungi on rumen volatile fatty acid concentrations in vivo. J Agri Sci 109:13–17.

10. Huws SA, Creevey CJ, Oyama LB, Mizrahi I, Denman SE, Popova M, Muñoz-Tamayo R, Forano E, Waters SM, Hess M, Tapio I, Smidt H, Krizsan SJ, Yáñez-Ruiz DR, Belanche A, Guan L, Gruninger RJ, McAllister TA, Newbold CJ, Roehe R, Dewhurst RJ, Snelling TJ, Watson M, Suen G, Hart EH, Kingston-Smith AH, Scollan ND, do Prado RM, Pilau EJ, Mantovani HC, Attwood GT, Edwards JE, McEwan NR, Morrisson S, Mayorga OL, Elliott C, Morgavi DP. 2018. Addressing global ruminant agricultural challenges through understanding the rumen microbiome: past, present, and future. Front Microbiol 9:2161.

11. Hanafy RA, Johnson B, Youssef NH, Elshahed MS. 2020. Assessing anaerobic gut fungal diversity in herbivores using D1/D2 large ribosomal subunit sequencing and multi-year isolation. Environ Microbiol 22:3883–3908.

12. Liggenstoffer AS, Youssef NH, Couger MB, Elshahed MS. 2010. Phylogenetic diversity and community structure of anaerobic gut fungi (phylum *Neocallimastigomycota*) in ruminant and non-ruminant herbivores. ISME J 4:1225–1235.

13. Meili CH, Jones AL, Arreola AX, Habel J, Pratt CJ, Hanafy RA, Wang Y, Yassin AS, TagElDein MA, Moon CD, Janssen PH, Shrestha M, Rajbhandari P, Nagler M, Vinzelj JM, Podmirseg SM, Stajich JE, Goetsch AL, Hayes J, Young D, Fliegerova K, Grilli DJ, Vodička R, Moniello G, Mattiello S, Kashef MT, Nagy YI, Edwards JA, Dagar SS, Foote AP, Youssef NH, Elshahed MS. 2023. Patterns and determinants of the global herbivorous mycobiome. Nat Commun 14:3798.

14. Young D, Joshi A, Huang L, Munk B, Wurzbacher C, Youssef NH, Elshahed MS, Moon CD, Ochsenreither K, Griffith GW, Callaghan TM, Sczyrba A, Lebuhn M, Flad V. 2022. Simultaneous metabarcoding and quantification of *Neocallimastigomycetes* from environmental samples: Insights into community composition and novel lineages. Microorganisms 10:1749.

15. Moon CD, Carvalho L, Kirk MR, McCulloch AF, Kittelmann S, Young W, Janssen PH, Leathwick DM. 2021. Effects of long-acting, broad spectra anthelmintic treatments on the rumen microbial community compositions of grazing sheep. Sci Rep 11:3836.

16. Azad E, Fehr KB, Derakhshani H, Forster R, Acharya S, Khafipour E, McGeough E, McAllister TA. 2020. Interrelationships of fiber-associated anaerobic fungi and bacterial communities in the rumen of bloated cattle grazing alfalfa. Microorganisms 8:1543.

17. Guo W, Wang W, Bi S, Long R, Ullah F, Shafiq M, Zhou M, Zhang Y. 2020. Characterization of anaerobic rumen fungal community composition in yak, Tibetan sheep and small tail Han sheep grazing on the Qinghai-Tibetan Plateau. Animals 10:144.

18. Kittelmann S, Seedorf H, Walters WA, Clemente JC, Knight R, Gordon JI, Janssen PH. 2013. Simultaneous amplicon sequencing to explore co-occurrence patterns of bacterial, archaeal and eukaryotic microorganisms in rumen microbial communities. PLOS ONE 8:e47879.

19. Kumar S, Indugu N, Vecchiarelli B, Pitta DW. 2015. Associative patterns among anaerobic fungi, methanogenic archaea, and bacterial communities in response to changes in diet and age in the rumen of dairy cows. Front Microbiol 6:781.

20. Hartinger T, Zebeli Q. 2021. The present role and new potentials of anaerobic fungi in ruminant nutrition. J Fungi (Basel) 7:200.

21. de Oliveira MN, Jewell KA, Freitas FS, Benjamin LA, Tótola MR, Borges AC, Moraes CA, Suen G. 2013. Characterizing the microbiota across the gastrointestinal tract of a Brazilian Nelore steer. Vet Microbiol 164:307–314.

22. Wang K, Zhang H, Hu L, Zhang G, Lu H, Luo H, Zhao S, Zhu H, Wang Y. 2022. Characterization of the microbial communities along the gastrointestinal tract in crossbred cattle. Animals (Basel) 12:825.

23. Swift CL, Louie KB, Bowen BP, Hooker CA, Solomon KV, Singan V, Daum C, Pennacchio CP, Barry K, Shutthanandan V, Evans JE, Grigoriev IV, Northen TR, O’Malley MA. 2021. Cocultivation of anaerobic fungi with rumen bacteria establishes an antagonistic relationship. mBio 12:e01442–21.

24. Henderson G, Cox F, Ganesh S, Jonker A, Young W, Abecia L, Angarita E, Aravena P, Nora Arenas G, Ariza C, Attwood GT, Mauricio Avila J, Avila-Stagno J, Bannink A, Barahona R, Batistotti M, Bertelsen MF, Brown-Kav A, Carvajal AM, Cersosimo L, Vieira Chaves A, Church J, Clipson N, Cobos-Peralta MA, Cookson AL, Cravero S, Cristobal Carballo O, Crosley K, Cruz G, Cerón Cucchi M, de la Barra R, De Menezes AB, Detmann E, Dieho K, Dijkstra J, dos Reis WLS, Dugan MER, Hadi Ebrahimi S, Eythórsdóttir E, Nde Fon F, Fraga M, Franco F, Friedeman C, Fukuma N, Gagić D, Gangnat I, Javier Grilli D, Guan LL, Heidarian Miri V, Hernandez-Sanabria E, et al. 2015. Rumen microbial community composition varies with diet and host, but a core microbiome is found across a wide geographical range. Sci Rep 5:14567.

25. Elshahed MS, Hanafy RA, Cheng Y, Dagar SS, Edwards JE, Flad V, Fliegerová KO, Griffith GW, Kittelmann S, Lebuhn M, O’Malley MA, Podmirseg SM, Solomon KV, Vinzelj J, Young D, Youssef NH. 2022. Characterization and rank assignment criteria for the anaerobic fungi (*Neocallimastigomycota*). Int J Syst Evol Microbiol 72: doi: 10.1099/ijsem.0.005449.

26. White JR, Nagarajan N, Pop M. 2009. Statistical methods for detecting differentially abundant features in clinical metagenomic samples. PLoS Comput Biol 5:e1000352.

27. Adrienne LJ, Carrie JP, Casey HM, Rochelle MS, Philip H, Mostafa SE, Noha HY. 2023. Anaerobic gut fungal communities in marsupial hosts. bioRxiv doi:10.1101/2023.05.31.543067:2023.05.31.543067.

28. Castillo C, Hernández J. 2021. Ruminal fistulation and cannulation: A necessary procedure for the advancement of biotechnological research in ruminants. Animals (Basel) 11:1870.

29. Hagey JV, Laabs M, Maga EA, DePeters EJ. 2022. Rumen sampling methods bias bacterial communities observed. PLoS One 17:e0258176.

30. Solden LM, Naas AE, Roux S, Daly RA, Collins WB, Nicora CD, Purvine SO, Hoyt DW, Schückel J, Jørgensen B, Willats W, Spalinger DE, Firkins JL, Lipton MS, Sullivan MB, Pope PB, Wrighton KC. 2018. Interspecies cross-feeding orchestrates carbon degradation in the rumen ecosystem. Nat Microbiol 3:1274–1284.

31. Liu JH, Zhang ML, Zhang RY, Zhu WY, Mao SY. 2016. Comparative studies of the composition of bacterial microbiota associated with the ruminal content, ruminal epithelium and in the faeces of lactating dairy cows. Microb Biotechnol 9:257–68.

32. Williamson JR, Callaway TR, Lourenco JM, Ryman VE. 2022. Characterization of rumen, fecal, and milk microbiota in lactating dairy cows. Front Microbiol 13:984119.

